# Biogenesis, engineering and function of membranes in the CO_2_-fixing pyrenoid

**DOI:** 10.1101/2024.08.08.603944

**Authors:** Jessica H. Hennacy, Nicky Atkinson, Angelo Kayser-Browne, Sabrina L. Ergun, Eric Franklin, Lianyong Wang, Moshe Kafri, Friedrich Fauser, Josep Vilarrasa-Blasi, Robert E. Jinkerson, Alistair J. McCormick, Martin C. Jonikas

## Abstract

Approximately one-third of global CO_2_ assimilation is performed by the pyrenoid^1^, a liquid-like organelle found in most algae and some plants^2^. Specialized membranes are hypothesized to drive CO_2_ assimilation in the pyrenoid by delivering concentrated CO_2_^3,4^, but their biogenesis and function have not been experimentally characterized. Here, we show that homologous proteins SAGA1 and MITH1 mediate the biogenesis of the pyrenoid membrane tubules in the model alga *Chlamydomonas reinhardtii* and are sufficient to reconstitute pyrenoid-traversing membranes in a heterologous system, the plant *Arabidopsis thaliana*. SAGA1 localizes to the regions where thylakoid membranes transition into tubules and is necessary to initiate tubule formation. MITH1 localizes to the tubules and is necessary for their extension through the pyrenoid. Tubule-deficient mutants exhibit growth defects under CO_2_-limiting conditions, providing evidence for the function of membrane tubules in CO_2_ delivery to the pyrenoid. Furthermore, these mutants form multiple aberrant condensates of pyrenoid matrix, indicating that a normal tubule network promotes the coalescence of a single pyrenoid. The reconstitution of pyrenoid-traversing membranes in a plant represents a key milestone toward engineering a functional pyrenoid into crops for improving crop yields. More broadly, our study demonstrates the functional importance of pyrenoid membranes, identifies key biogenesis factors, and paves the way for the molecular characterization of pyrenoid membranes across the tree of life.

## Main Text

The growth of photosynthetic organisms is commonly limited by the rate of CO_2_ fixation catalyzed by the enzyme Rubisco^5^. Nearly all eukaryotic algae and some plants overcome this limitation with a chloroplast-localized organelle called the pyrenoid, which enhances CO_2_ fixation (Fig. 1a)^2^. The prevalence of pyrenoids in eukaryotic algae has led to the estimate that pyrenoids mediate 61-88% of primary productivity in the oceans^1^, making the pyrenoid an organelle of major biogeochemical importance. Furthermore, engineering a pyrenoid into major C3 crops such as wheat and rice is expected to enhance their CO_2_ uptake, yield, and water use efficiency^6^, which will promote agricultural resilience in the face of climate change^7^.

**Fig. 1.**
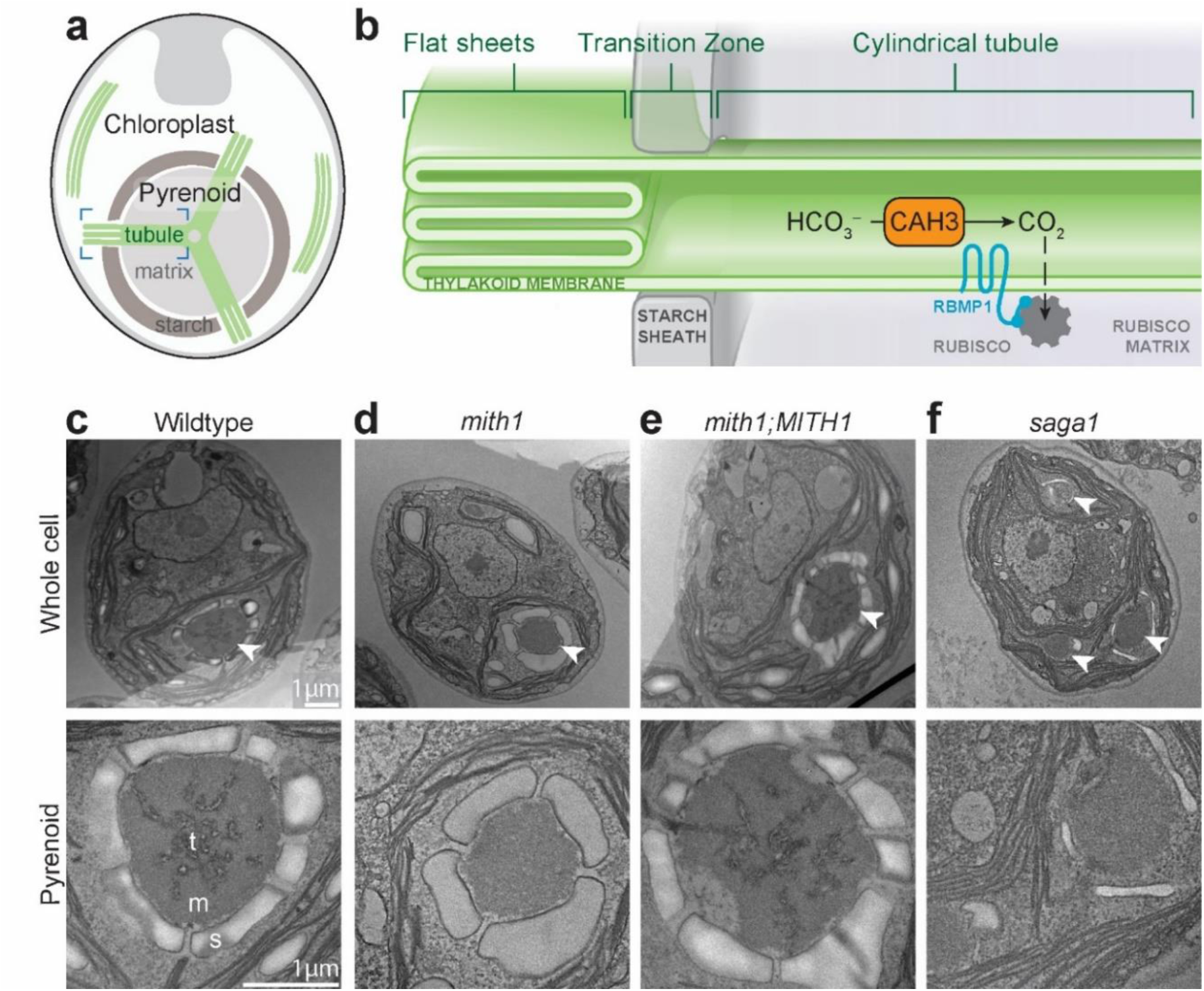
MITH1 and SAGA1 are required for pyrenoid-traversing thylakoid membrane tubules. **a**, The Chlamydomonas pyrenoid is found at the base of the cell’s cup-shaped chloroplast and consists of a Rubisco matrix traversed by membrane tubules and surrounded by a starch sheath. **b**, Flat sheets of thylakoid membrane transition into cylindrical tubules as they enter the pyrenoid matrix through gaps in the starch sheath. Bicarbonate (HCO_3_^-^) is concentrated in the tubule lumen and converted to CO_2_ by CAH3, Carbonic Anhydrase 3^25^. RBMP1, Rubisco Binding Membrane Protein 1^18^, is a putative tubule-matrix linker protein. **c-f**, Transmission electron microscopy (TEM) images of whole cells and zoom-in on each cell’s pyrenoid. Arrowheads point to pyrenoids. **c**, t: tubule, m: matrix, s: starch. **e**, In the *mith1;MITH1* strain, the tagged protein MITH1-Venus-3xFLAG is expressed in a *mith1* mutant background.

Pyrenoids enhance CO_2_ fixation by supplying Rubisco with concentrated CO_2_. All known pyrenoids share two structural features: a spheroidal protein matrix and specialized membranes (Fig. 1a)^2^. The matrix is the site of CO_2_ fixation by Rubisco, and its biogenesis has recently been elucidated in the model alga *Chlamydomonas reinhardtii* (Chlamydomonas hereafter); it is a liquid-like condensate^8^ composed primarily of Rubisco and an intrinsically disordered linker repeat protein, EPYC1^1,9,10^.

Pyrenoid-associated membranes have been hypothesized to serve the key function of supplying concentrated CO_2_ to the matrix (Fig. 1b)^11^. By contrast with the matrix, the biogenesis and function of these membranes have not been experimentally characterized in any organism due to the lack of any mutant known to directly disrupt them.

### MITH1 and SAGA1 are necessary for pyrenoid tubule formation

A high-throughput mutant screen in Chlamydomonas recently identified *MITH1* (*MIssing THylakoids 1, Cre06.g259100*, *SAGA3*) as a gene required for normal growth in low CO_2_, a characteristic of pyrenoid-related genes^12^. In the present study, using transmission electron microscopy (TEM), we found that normal pyrenoid membranes were absent in *mith1* mutants (Fig. 1 and Extended Data Fig. 1). In wildtype Chlamydomonas, pyrenoid membranes appear as cylindrical tubules that traverse the matrix and merge with the photosynthetic thylakoid membrane sheets outside the pyrenoid (Fig. 1a-c). In the *mith1* mutant, traversing tubules were absent from the pyrenoid matrix (Fig. 1d). To our knowledge, this is the first observation of a mutant with a phenotype specifically affecting the tubules. In the complemented strain *mith1*; *MITH1-Venus*, a normal pyrenoid-traversing tubule network is restored (Fig. 1e), demonstrating that MITH1 is necessary for the formation of pyrenoid-traversing tubules.

The discovery of a role for MITH1 in pyrenoid membrane tubule formation suggested a similar function for its homolog, SAGA1^12^, a protein that is also required for growth under low CO_2_ but whose function had remained elusive due to the *saga1* mutant’s complex phenotypes that impacted multiple pyrenoid sub-compartments^13^. The idea that SAGA1 functions in the same tubule biogenesis pathway as MITH1 is supported by the observation that *mith1* and *saga1* mutants displayed a similar pattern of growth defects across 121 different conditions in a high-throughput study^12^. Further consistent with a possible function for SAGA1 in tubule biogenesis, only ∼8% of *saga1* pyrenoid matrix condensates contained visible membranes, and these membranes showed abnormal morphology (Fig. 1f)^13^.

### MITH1 and SAGA1 are sufficient to produce matrix-traversing membranes

To determine if MITH1 and SAGA1 are sufficient for making pyrenoid matrix-traversing membranes, we expressed the two proteins in a heterologous system: *Arabidopsis thaliana* plants (Arabidopsis hereafter) (Fig. 2, Extended Data Fig. 2). In this system, a synthetic Chlamydomonas-like pyrenoid matrix was previously reconstituted by expressing the linker protein EPYC1 alongside the EPYC1-binding Chlamydomonas Rubisco small subunit CrRBCS2. While wild-type Arabidopsis chloroplasts lack a pyrenoid (Fig. 2a), expression of EPYC1-GFP and CrRBCS2 was sufficient for sequestering Rubisco in a single large condensate within the chloroplast^14^ (Fig. 2b,f). Here, we generated plants that express SAGA1-mCherry and chloroplast-targeted MITH1-mCerulean in addition to EPYC1-GFP and CrRBCS2 (See Methods). SAGA1-mCherry and MITH1-mCerulean both localized to the CrRBCS2;EPYC1-GFP matrix condensates in these plants (Fig. 2d,e, Extended Data Fig. 3), indicating that once in the chloroplast, both proteins are targeted to the matrix.

**Fig. 2.**
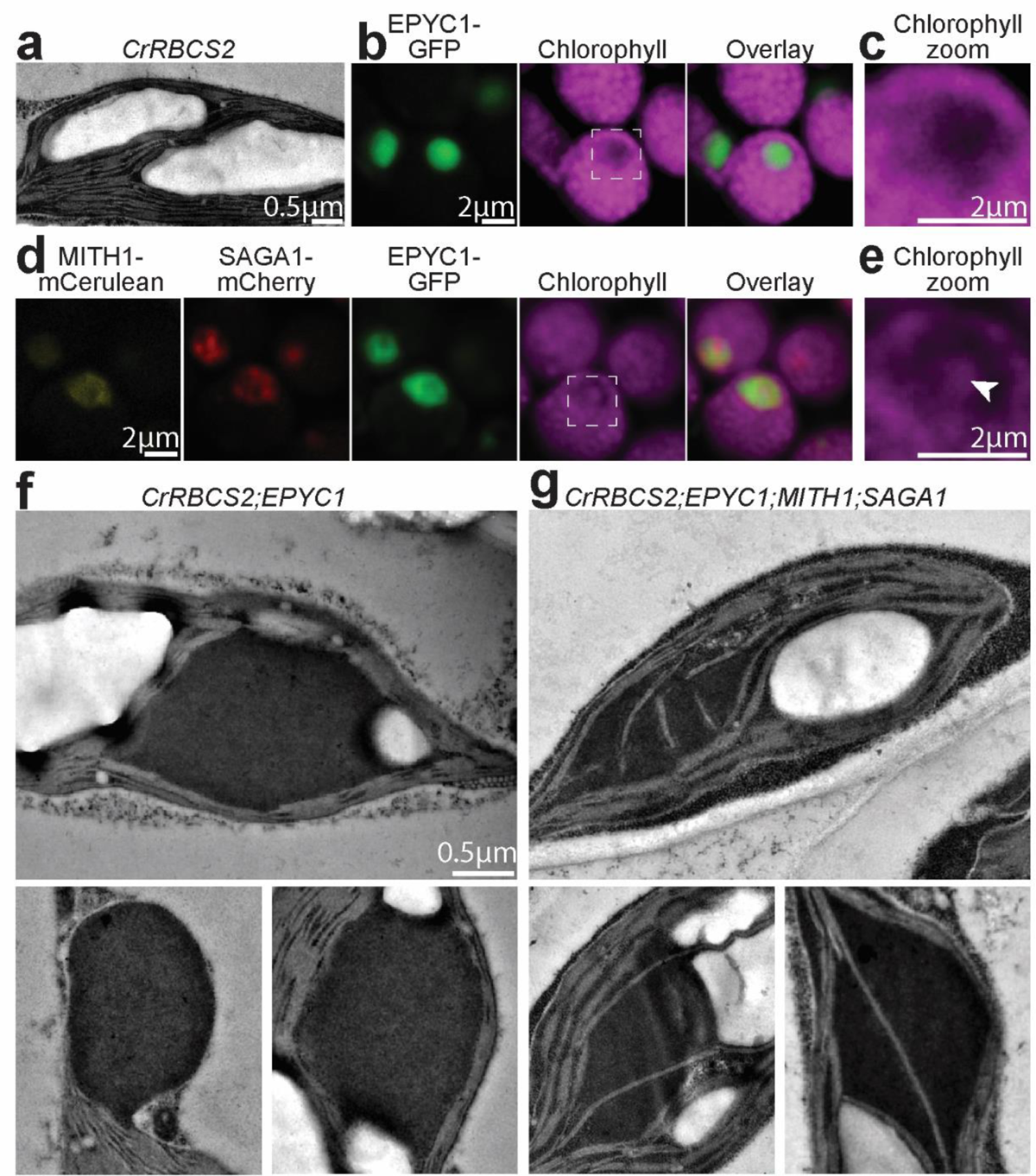
MITH1 and SAGA1 are sufficient to form Rubisco matrix-traversing thylakoid membranes in *Arabidopsis*. **a**, TEM image of a chloroplast from the Arabidopsis CrRBCS2 line, where a Chlamydomonas Rubisco small subunit (CrRBCS2) is expressed^26^. **b**, Confocal images depicting the *CrRBCS2*;*EPYC1-GFP* line^14^. **c**, Zoom-in on chlorophyll in the region of the CrRBCS2;EPYC1-GFP condensate, as indicated in (**b**). **d**, Confocal microscopy images depicting the *Arabidopsis* line *CrRBCS2;EPYC1-GFP;MITH1-mCerulean;SAGA1-mCherry*. **e**, Zoom-in on the chlorophyll autofluorescence channel in the region indicated in panel (**d)**. Arrowhead points to chlorophyll signal inside the CrRBCS2;EPYC1-GFP condensate. **e, f** TEM images of condensates in the original *CrRBCS2;EPYC1-GFP* line^14^ (**e**) alongside the *CrRBCS2;EPYC1-GFP;MITH1-mCerulean;SAGA1-mCherry* line (**f**).

Whereas membranes had never previously been observed within condensates of the *CrRBCS2;EPYC1-GFP* Arabidopsis line (Fig. 2c), we detected chlorophyll fluorescence puncta inside the condensates of the *CrRBCS2;EPYC1-GFP;MITH1-mCerulean;SAGA1-mCherry* line, suggesting that they contain thylakoid membranes (Fig. 2e). TEM confirmed that thylakoid membranes traverse the matrix condensates in this line (Fig. 2g); we observed traversing membranes in 32% of *CrRBCS2;EPYC1-GFP;MITH1-mCerulean;SAGA1-mCherry* condensates (n=166), but in none of the condensates of the control line (n=117; p < 0.0001, Fisher’s exact test; Fig. 2f). These results demonstrate that MITH1 and SAGA1 are sufficient to generate matrix-traversing thylakoid membranes in a heterologous system. The discovery of proteins that are both necessary and sufficient for formation of matrix-traversing membranes now enables investigation of the process of pyrenoid membrane biogenesis and their functions.

### MITH1 and SAGA1 function at distinct regions of the tubules

Whereas MITH1 was previously proposed to localize to the pyrenoid matrix^15,16^, here we demonstrate that it localizes and functions in the pyrenoid tubules. We examined the localization of MITH1-Venus in the complemented *mith1;MITH1-Venus* background where the tagged protein is known to be functional (Fig. 1e). MITH1 localized to a branching pattern in the pyrenoid similar to that of RBMP1, a known tubule protein, and distinct from the homogeneous localization of the Rubisco small subunit (RBCS1), which is known to occupy the matrix (Fig. 3a-c). Further supporting tubule localization, cell fractionation revealed that MITH1 co-pelleted with membranes in wild-type (Extended Data Fig. 4) and in a matrix-less mutant (Fig. 3h). Together, these results demonstrate that MITH1 localizes to the pyrenoid tubules.

**Fig. 3.**
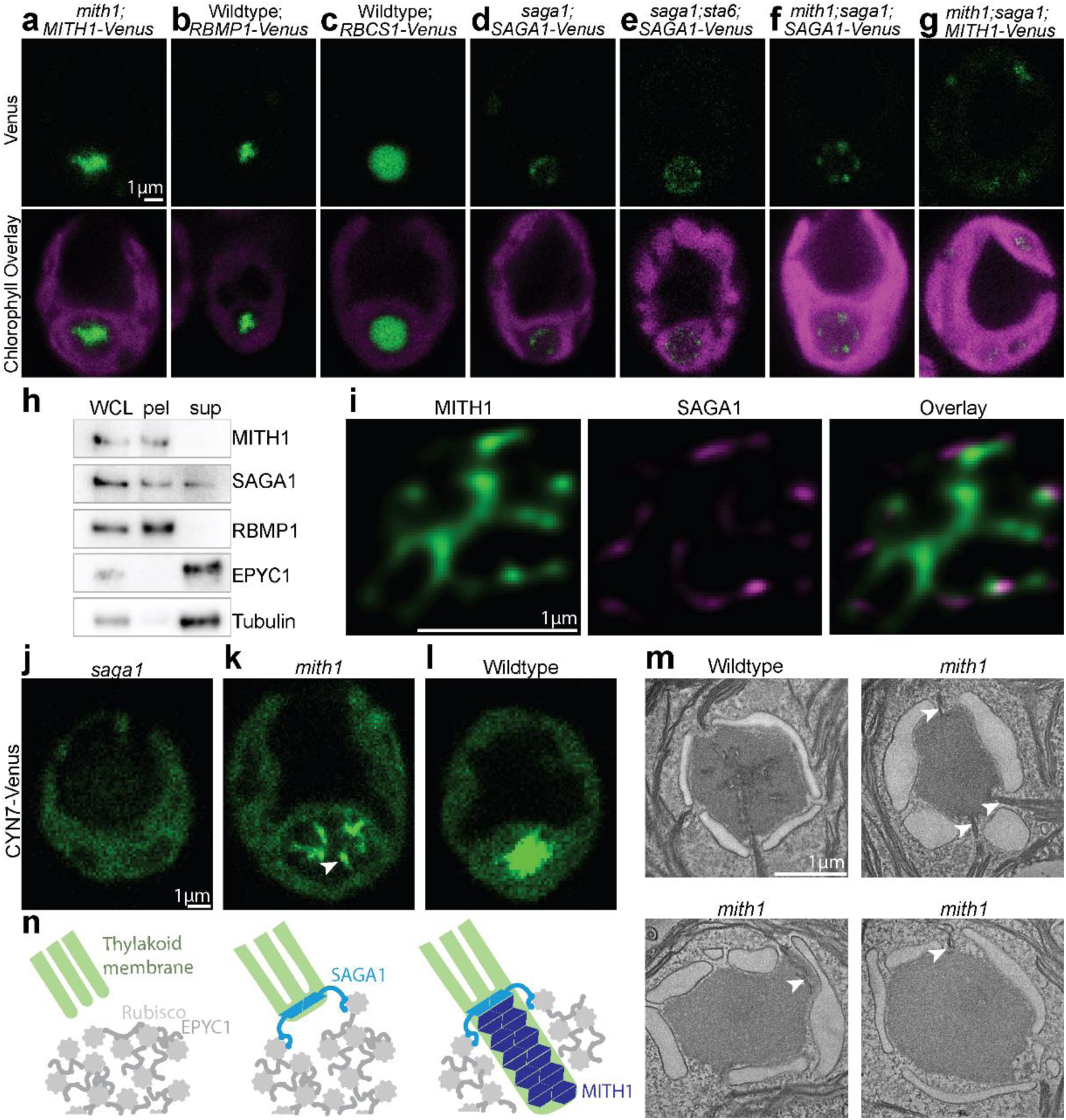
SAGA1 and MITH1 function sequentially at distinct regions of the tubules. **a-g**, Localizations of Venus-tagged proteins in various mutant backgrounds. **h,** Cell fractionation and western blot of the matrix-less D/E mutant^10^. **i,** STimulated Emission Depletion (STED) super-resolution microscopy of MITH1 and SAGA1-Venus-3xFLAG with anti-MITH1 and anti-FLAG immunofluorescence. **j-l,** Localization of CYN7-Venus in wildtype versus mutant backgrounds. **m,** TEM of wildtype and *mith1* mutant pyrenoids. Arrowheads point to thylakoid protrusions. **n,** Model: SAGA1 initiates tubule biogenesis from thylakoid membranes and mediates contact between membranes and Rubisco matrix; MITH1 extends tubules through the matrix.

We had previously found that SAGA1-Venus localizes to puncta at the pyrenoid periphery and had hypothesized that this localization was mediated by its CBM20 starch-binding domain^13^, which could bind the starch sheath that surrounds the Chlamydomonas pyrenoid (Fig. 1a). Here, we found that SAGA1 localizes to pyrenoid-associated puncta even in the starch-less *sta6* mutant background (Fig. 3e), indicating that SAGA1 does not require starch for its normal localization and raising the possibility that it is targeted to the membranes instead. We detected SAGA1 both in the membrane pellet and supernatant in the cell fractionation experiment, indicating that while SAGA1 interacts with membranes, a portion of the protein dissociates from membranes during the experiment or exists in a distinct soluble state (Fig. 3h).

To examine the co-localization of MITH1 and SAGA1 we used Stimulated Emission Depletion (STED) super-resolution microscopy (Fig. 3i and Extended Data Fig. 5). SAGA1 localized to puncta at the extremities of MITH1’s branched localization pattern. These results suggest that SAGA1 and MITH1 localize to different topological regions of the tubules, with MITH1 on the cylindrical portion of tubules and SAGA1 at the transition zones at the tubule-thylakoid membrane interfaces at the surface of the pyrenoid.

### MITH1 requires SAGA1 for localization and function

The localizations of MITH1 and SAGA1 to distinct regions of the tubules suggest that MITH1 and SAGA1 act at different steps of the tubule biogenesis pathway. To determine the order in which MITH1 and SAGA1 act in tubule biogenesis, we tested their epistasis by generating a *mith1;saga1* double mutant. We compared the growth and cell morphology of single and double mutants using spot tests, light microscopy, and TEM. The *mith1*;*saga1* double mutant displayed similar growth and morphological defects to the *saga1* single mutant (Extended Data Fig. 6), demonstrating that *saga1* is epistatic to *mith1*, and suggesting that SAGA1 function is necessary for MITH1 function.

Furthermore, we found that SAGA1 localized correctly in the absence of MITH1, but not vice-versa. We expressed each Venus-tagged protein in a *mith1;saga1* double mutant (Fig. 3f,g). SAGA1-Venus showed its normal localization to pyrenoid puncta in this background, where MITH1 is not expressed (Fig. 3f, Extended Data Fig. 7). In contrast, MITH1-Venus mis-localized throughout the chloroplast in this background, where SAGA1 is absent (Fig. 3g, Extended Data Fig. 7). Together with the epistasis results above, these findings demonstrate that SAGA1 can localize and function independently of MITH1, whereas MITH1 requires SAGA1 for its normal localization and function.

### SAGA1 initiates tubules and MITH1 extends them

Our results so far have shown that SAGA1 and MITH1 function at different sites and stages of tubule biogenesis. To distinguish between the specific roles of SAGA1 and MITH1, we characterized tubule defects in *saga1* and *mith1* using the protein CYN7-Venus as a marker for tubules. Whereas CYN7-Venus was clearly enriched in the tubules of wildtype cells (Fig. 3l)^17^, in the *saga1* mutant CYN7-Venus was dispersed throughout the chloroplast (Fig. 3j), demonstrating that a normal tubule network is not present anywhere in cells of this mutant and indicating that SAGA1 is required for tubule initiation at the cup of the chloroplast. Unlike in the *saga1* mutant, in the *mith1* mutant CYN7-Venus localized to streaks and puncta associated with the periphery of the pyrenoid (Fig. 3k and Extended Data Fig. 8).

Consistent with these observations, several TEM images of the *mith1* mutant showed thylakoid membranes crossing the pyrenoid starch sheath and stopping at the surface of the matrix (Fig. 3m, arrowhead). These observations indicate that tubule formation initiates at the correct cellular location in the *mith1* mutant but then stalls before tubules are extended through the pyrenoid matrix. SAGA1’s punctate localization in *mith1* mutants (Fig. 3f) suggests that SAGA1 is present in these streaks and functions to form them.

Based on our results, we propose a model in which SAGA1 initiates tubule biogenesis at the periphery of the pyrenoid, while MITH1 extends tubules through the matrix (Fig. 3n). We have previously observed that SAGA1 binds to Rubisco in a yeast two-hybrid assay^13^ and more recently found that SAGA1 contains a Rubisco-binding motif that binds to Rubisco in vitro^18^. Thus, in our model, we propose that SAGA1 initiates contact between membranes and the matrix via its Rubisco-binding motifs. MITH1 then promotes growth of the tubule inward through the matrix.

### Tubule-deficient mutants have multiple pyrenoid matrix condensates

Our discovery of tubule-deficient mutants now provides an opportunity to gain insights into the impacts that these pyrenoid-associated membranes have on the overall structure and function of the pyrenoid. We had previously observed that the *saga1* mutant formed an average of 10 matrix condensates throughout the chloroplast, and had proposed that this was a consequence of starch defects (Fig. 4a)^13^. Our discovery that SAGA1 mediates tubule biogenesis now leads us to hypothesize that the multiple matrix condensates in the *saga1* mutant are instead a consequence of tubule defects.

**Fig. 4.**
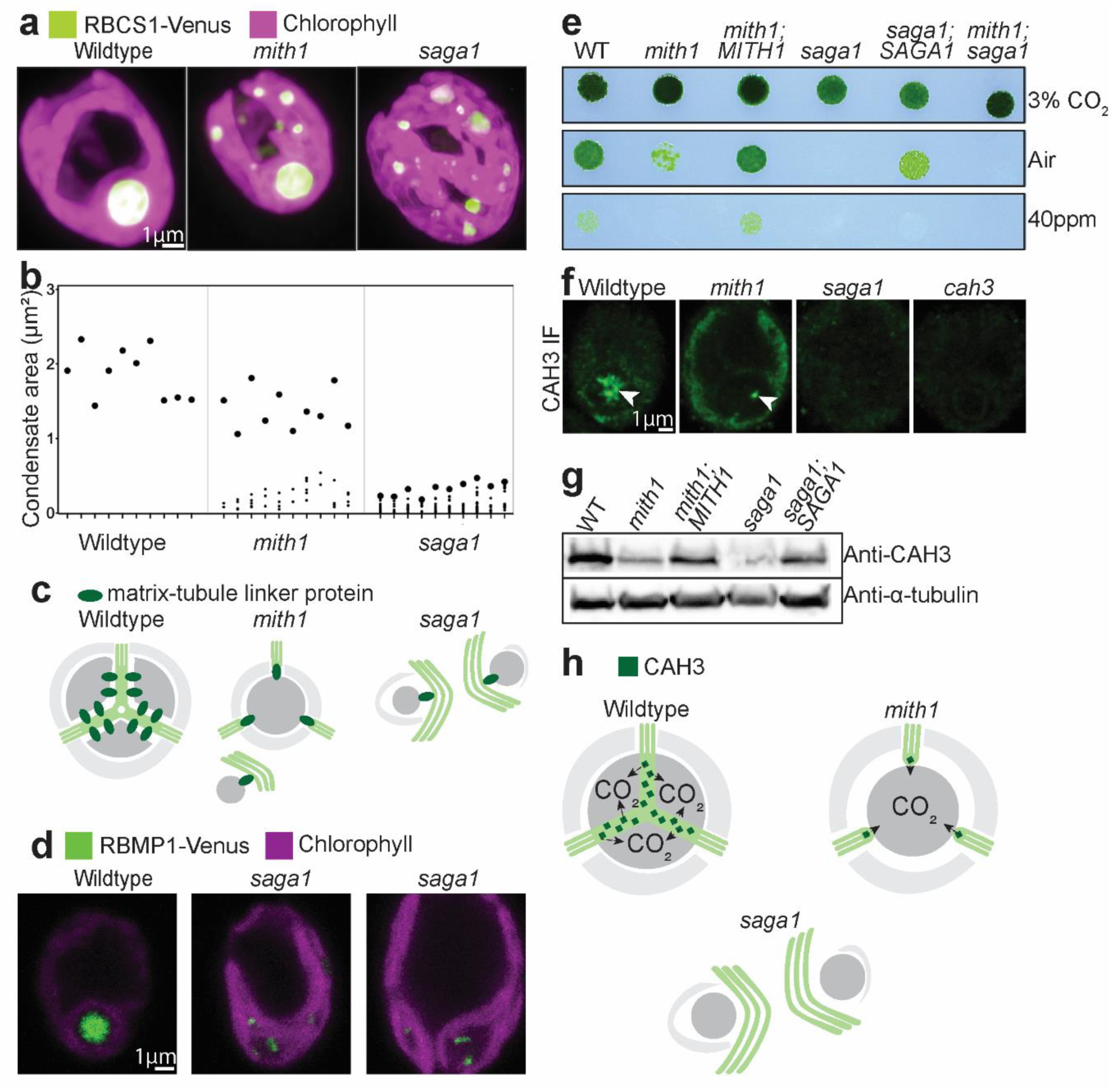
Tubule-deficient mutants have multiple pyrenoids and defective CO_2_ delivery to the pyrenoid. **a**, Max-z projections depicting RBCS1-Venus localization in wildtype, *mith1*, and *saga1* cells. The chloroplast is visualized by chlorophyll autofluorescence. **b**, Graph depicting the largest cross-sectional area observed in any z-slice for each condensate in a given cell. Each tick on the x-axis corresponds to a single cell; each point on the graph plots the area for a single condensate. The point depicting the largest condensate in each cell is enlarged to aid visualization. Data for 10 cells per strain are shown. **c**, Model for how mis-localized tubule-matrix linker proteins could drive mis-localization of Rubisco in *mith1* and *saga1* mutants. In the absence of tubules, mis-localized tubule-matrix linker proteins could nucleate Rubisco puncta throughout the chloroplast. **d,** Localization of putative matrix-tubule linker RBMP1 in wildtype versus *saga1* mutant backgrounds. **e,** Spot test depicting growth at varying levels of CO_2_ and 100 µmol photons m^−2^⋅s^−1^ light levels. **f,** Anti-CAH3 immunofluorescence. Arrowheads point to regions of strong CAH3 signal associated with the pyrenoid. **g**, Anti-CAH3 Western blot performed on whole cell lysates form different samples. Anti-alpha-tubulin is used as a loading control. **h,** Model: Partially localized CAH3 provides enough CO_2_ to sustain partial growth in the *mith1* mutant.

The idea that tubule biogenesis defects lead to multiple matrix condensates is supported by two observations. First, we disproved the major alternative hypothesis that the multiple matrix condensates in *saga1* are caused by the abnormal starch in this mutant^13^ by observing that the starch-less *saga1;sta6* double mutant still forms multiple matrix condensates (Extended Data Fig. 9). Second, the *mith1* mutant also exhibited multiple matrix condensates (Fig. 4a). The *mith1* phenotype was less pronounced, with a single major pyrenoid present and fewer aberrant condensates (Fig. 4b). Considering that *mith1* has less-severe tubule defects than *saga1*, our observations indicate that the extent of the multiple-matrix-condensates phenotype is correlated with the extent of tubule biogenesis defects (Fig. 3j-l, 4a-b).

The observation that the Arabidopsis *CrRBCS2;EPYC1-GFP* plants contained only one condensate per chloroplast (Fig. 3b) even though they lack SAGA1 and MITH1 suggests that the multiple condensates in the Chlamydomonas *saga1* and *mith1* mutants are caused by the presence of one or more additional factors that are not present in the Arabidopsis CrRBCS2;EPYC1 plants. We therefore hypothesize that the multiple matrix condensates found throughout the chloroplast of the Chlamydomonas mutants are nucleated by mis-localized tubule-matrix linker proteins^18^.

In wildtype cells, the matrix is thought to be connected to tubules by tubule membrane-localized proteins that contain Rubisco-binding motifs^18^. We propose that in the *saga1* mutant, which lacks tubules, these tubule-matrix linker proteins mis-localize along thylakoid membrane sheets and nucleate matrix condensates throughout the chloroplast (Fig. 4c). By contrast, in the *mith1* mutant, which initiates tubule biogenesis, tubule-matrix linker proteins at the nascent tubules recruit most of the matrix to a major pyrenoid. Consistent with this hypothesis, in the *saga1* mutant, tubule-matrix linker RBMP1 mis-localized to multiple crescents that appeared to surround condensates throughout the chloroplast (Fig. 4d). These results reveal a previously-unappreciated role for tubules in establishing a single pyrenoid condensate.

### Tubules enable carbonic anhydrase accumulation and growth under limiting CO_2_

Tubules have long been proposed to deliver concentrated CO_2_ to the pyrenoid, however their functional importance in CO_2_ delivery had not been experimentally established due to the lack of mutants known to specifically affect them. Our discovery that *mith1* and *saga1* are deficient in tubule biogenesis now provides such mutants, allowing us to examine the consequences of tubule defects on pyrenoid function. These mutants exhibit growth defects in CO_2_-limiting conditions that are rescued by elevated CO_2_ (Fig. 4e)^12,13^, indicating that Rubisco in their pyrenoids is not supplied with sufficient CO_2_ to support growth in CO_2_-limiting conditions. Together, these observations provide experimental support for the proposed role of tubules in delivering concentrated CO_2_ to the pyrenoid.

Unexpectedly, we found that the *mith1* mutant’s growth defect is less severe than *saga1*’s at air levels of CO_2_ (Fig. 4e). A potential explanation for this difference is that the partially-initiated tubules found at the surface of *mith1* mutant pyrenoids supply enough CO_2_ to sustain growth in air. To examine CO_2_ delivery to the *mith1* pyrenoid, we determined the localization of the carbonic anhydrase CAH3 in this mutant. According to current models, CAH3 in the tubule lumen generates CO_2_ from a high concentration of bicarbonate (HCO_3_^-^)^2,3^. If the partially-initiated tubules seen in *mith1* mutants have any capacity to supply CO_2_, we would expect CAH3 to be found there. Indeed, we observed that CAH3 localized to pyrenoid-peripheral puncta in the *mith1* mutants (Fig. 4f) which likely correspond to the membrane protrusions seen by TEM (Fig. 3m). By contrast, in the *saga1* mutant, we could not detect CAH3 above background levels anywhere in the cell, indicating that *saga1* mutant pyrenoids lack a source of CO_2_.

We found that CAH3 protein levels are decreased in both mutants, but to a more severe extent in *saga1* (Fig. 4g). Furthermore, a recent study^19^ found that expression levels of other genes involved in carbon uptake, including HLA3 and LCIA, are decreased in the *saga1* mutant. Taken together, these observations indicate that tubules are required for accumulation of proteins involved in CO_2_ and HCO_3_^-^ transport. We propose that this regulatory requirement avoids the unproductive and energetically wasteful operation of the CO_2_-concentrating mechanism when tubules are not properly assembled and thus there is no effective route for delivery of CO_2_ to Rubisco. Our findings underscore the importance of tubules for CO_2_ delivery and reveal the existence of a regulatory mechanism that ensures that concentrated CO_2_ is only released by properly-assembled pyrenoid tubules.

### Perspective

Our findings elucidate the biogenesis and functions of membranes in the pyrenoid, an organelle that enhances photosynthetic CO_2_ assimilation in the oceans. We identify the first pyrenoid membrane biogenesis factors, determine how they assemble the pyrenoid tubules, and demonstrate the necessity of pyrenoid membranes for CO_2_ delivery. Traversing membranes are observed in nearly all pyrenoids^20^, but they have not been characterized at a molecular level in any other species. While the protein components may differ in other lineages due to the recent convergent evolution of pyrenoids, our work lays a blueprint for characterization of pyrenoid-traversing membrane biogenesis across the phylogenetic tree and reveals basic principles that we propose apply broadly.

Our discovery of pyrenoid membrane biogenesis factors allowed us to reconstitute pyrenoid-traversing membranes in a land plant, overcoming a major roadblock to pyrenoid engineering for increased yields. After the reconstitution of a Rubisco matrix in plants^14^, the reconstitution of tubules was widely seen as the key next step^3,20–22^, but progress on this front was stalled due to a complete lack of understanding of how the pyrenoid tubules are formed. Using the knowledge gained here, we have now produced thylakoid membranes that traverse an EPYC1-Rubisco matrix in *Arabidopsis*, achieving the reconstitution of all key structural features that we expect will be needed for a functional pyrenoid in a plant^3^. The plants we produced here are a powerful platform for testing additional pyrenoid biogenesis components and advancing efforts to install CO_2_ delivery pathways into these membranes over the coming decade.

Pyrenoid membrane tubules have been hypothesized to localize the pyrenoid to the base of the chloroplast cup, as the tubules still form in the canonical location even in the absence of matrix or starch^10,23^. Our observation that *mith1* still forms a major pyrenoid at the canonical location indicates that the central portion of the tubules is not needed for pyrenoid localization. However, *saga1* fails to localize any of the pyrenoid’s three sub-compartments to the base of the chloroplast, indicating that in addition to its role in tubule biogenesis, SAGA1 plays a key role in positioning the pyrenoid within the cell.

During cell division, multiple matrix condensates transiently appear throughout the chloroplast^8^. Our finding that the disruption of tubules leads to multiple matrix condensates leads us to hypothesize that the multiple condensates observed during cell division are due to transient disassembly of tubules and the resulting re-localization of tubule-matrix linkers. Such a process could contribute to the even partitioning of tubule proteins and matrix between daughter cells.

Membranes such as the nuclear envelope and plasma membrane have been proposed to play a role in promoting phase separation by scaffolding condensate components^24^. However, it has not been feasible to knock out these membrane structures *in vivo* to examine their impact on phase separation. Here, we have disrupted the membrane network central to the liquid-like pyrenoid and observed that this causes a single condensate to break into many, contributing evidence that membranes regulate the number and localization of condensates.

In conclusion, our molecular characterization of the pyrenoid tubules opens the door to their use as a model for investigating how membranes nucleate phase-separated condensates, their engineering toward enhanced crop yields, and their functional study toward understanding the biological mechanisms that underlie the global carbon cycle.

## Extended Data

**Extended Data Fig. 1.**
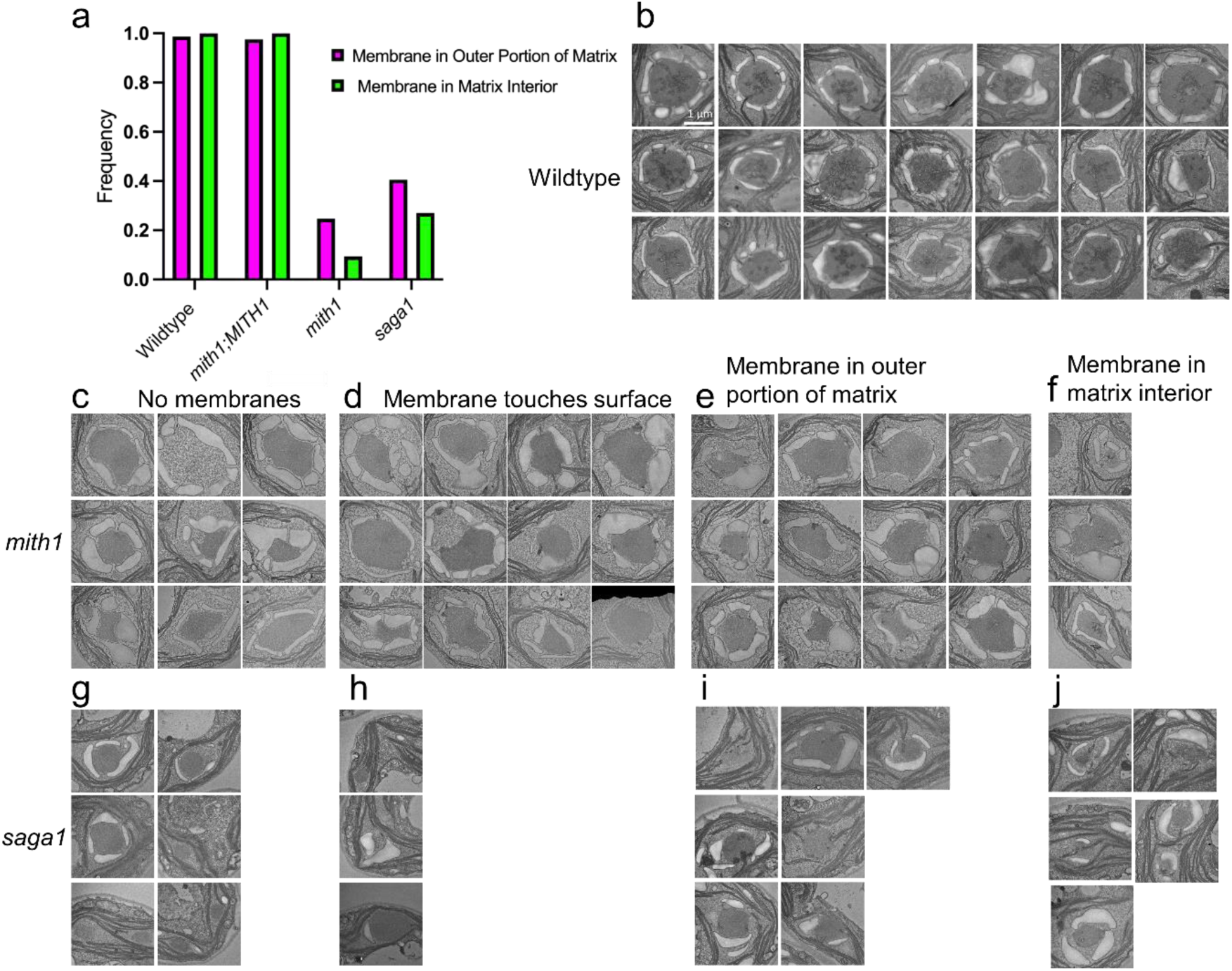
By TEM, *mith1* and *saga1* mutant pyrenoids are missing the canonical tubule network observed in wildtype pyrenoids. a, Graph depicting for each strain the percentage of pyrenoids with membranes seen entering the matrix surface or interior. **b**, TEM images of wildtype pyrenoids. **c-j**, TEM images of *mith1* and *saga1* mutant pyrenoids, organized into categories based on membrane contact with the pyrenoid. The “No membranes” (**c**, **g**) category indicates that no membranes are seen in the pyrenoid. The “Membrane touches surface” (**d**, **h**) category indicates that membranes are seen touching the surface of the pyrenoid, but are not seen entering the pyrenoid matrix. The “Membrane in outer portion of matrix” category (**e**, **i**) describes pyrenoids where membranes are seen entering the pyrenoid matrix at the surface. The “Membrane in matrix interior” category (**f**, **j**) includes pyrenoids where membrane is seen at the interior of the matrix.

**Extended Data Fig. 2.**
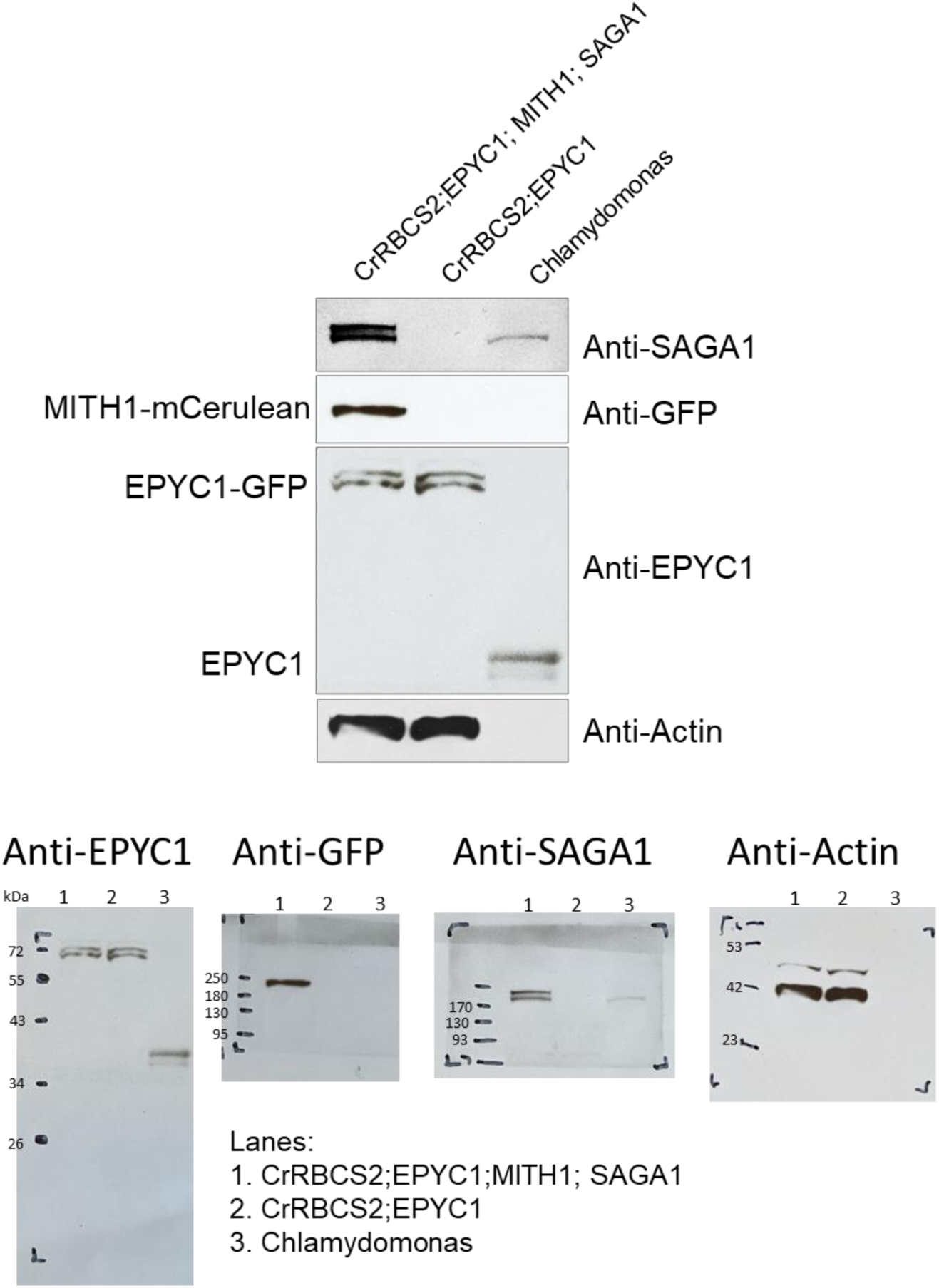
MITH1-mCerulean and SAGA1-mCherry were heterologously expressed in an *Arabidopsis* line containing CrRBCS1 and EPYC1-GFP. Western blots were used to probe for proteins expressed heterologously in *Arabidopsis* lines as compared to the native proteins from Chlamydomonas. The anti-GFP antibody was used to detect mCerulean. The bottom row of blots includes molecular weight markers.

**Extended Data Fig. 3.**
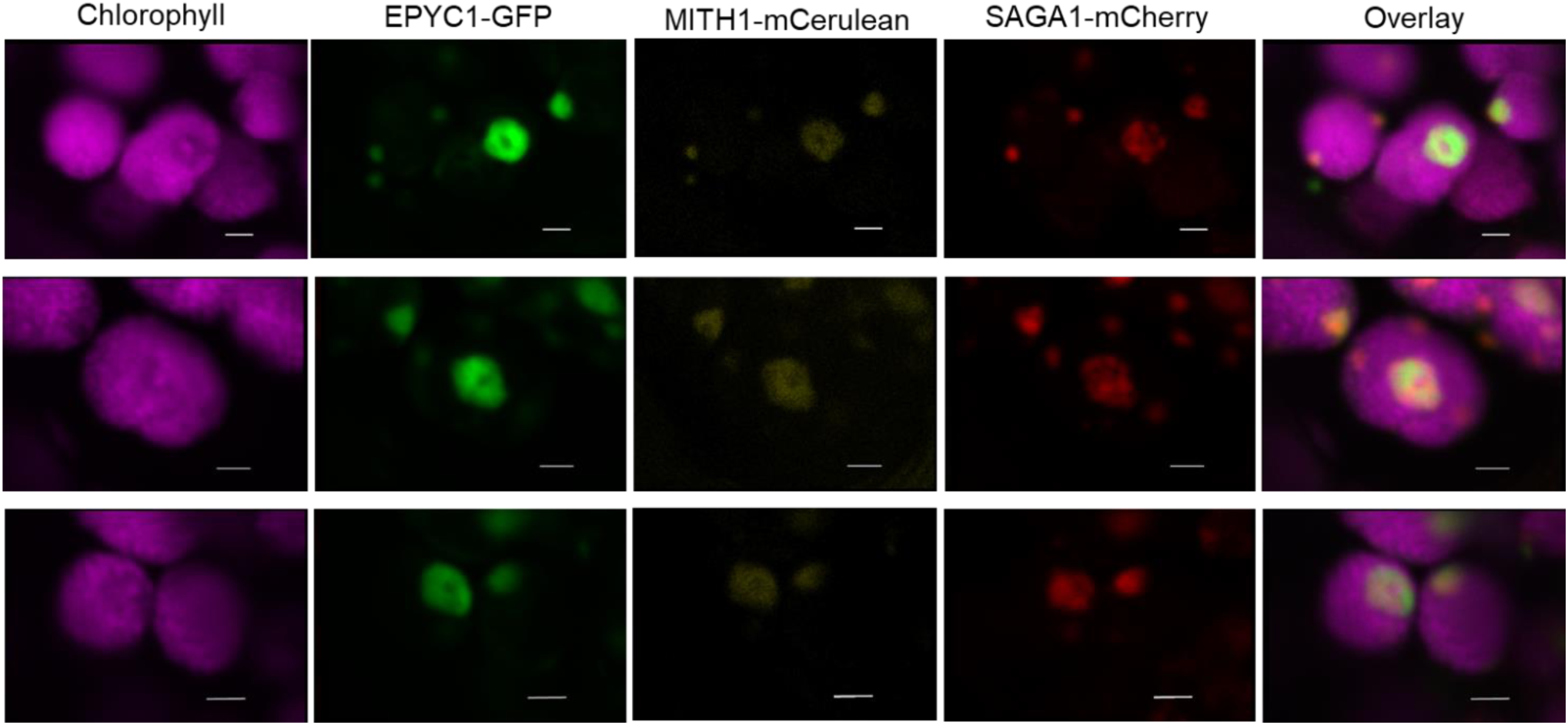
MITH1-mCerulean and SAGA1-mCherry co-localize with EPYC1-GFP. Additional images of heterologous protein localizations in *Arabidopsis* are shown. Scale bar: 2 μm.

**Extended Data Fig. 4.**
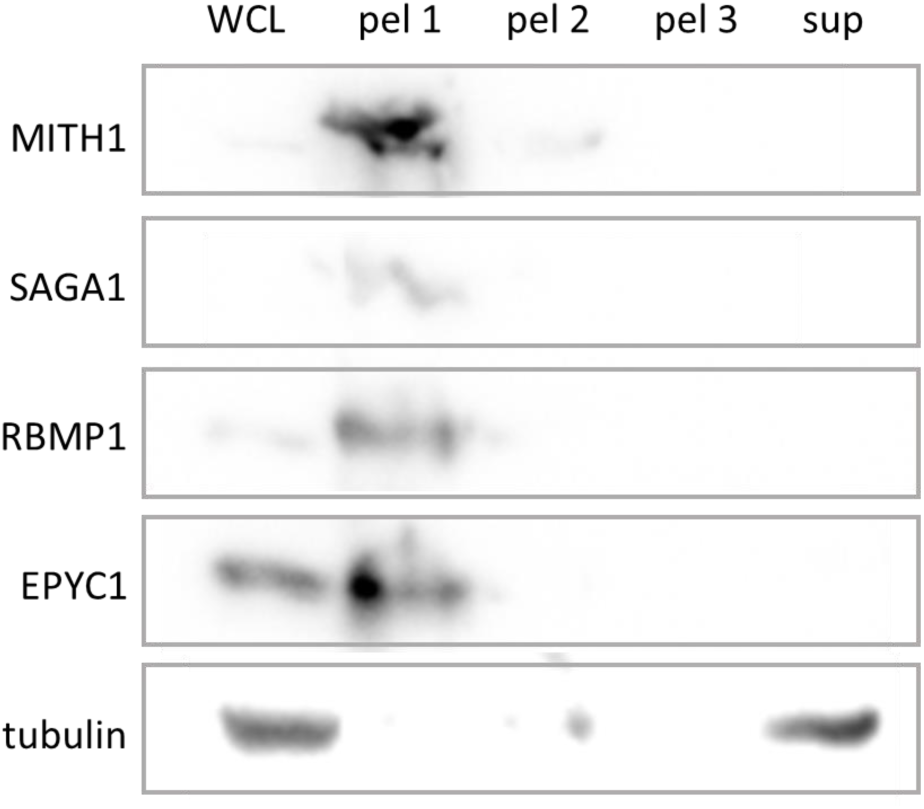
Using wildtype cells, the Rubisco matrix pellets with membranes in a cell fractionation experiment. Whole cell lysates (WCL) of the wildtype CMJ030 were sequentially centrifuged at 15,000 g (pel 1), 40,000 g (pel 2), and 70,000 g (pel 3). After each centrifugation step, the pellet (pel) was collected and the supernatant was used for the subsequent centrifugation step. The WCL, pellets (pel 1-3), and supernatant from the third centrifugation step (sup, concentrated 2x using a 10KDa centrifugal filter) were probed by western blot for the indicated proteins. The SAGA1 antibody cross-reacts with RBMP1 and EPYC1^18^, and these cross-reacting bands are shown here. RBMP1^18^ was used as a marker for tubules. EPYC1^1^, Essential Pyrenoid Component 1, links Rubisco in the matrix of wildtype cells, and was found in the pellet here, indicating that our centrifugation protocol does not separate the pyrenoid matrix from membranes when a wildtype strain is used.

**Extended Data Fig. 5.**
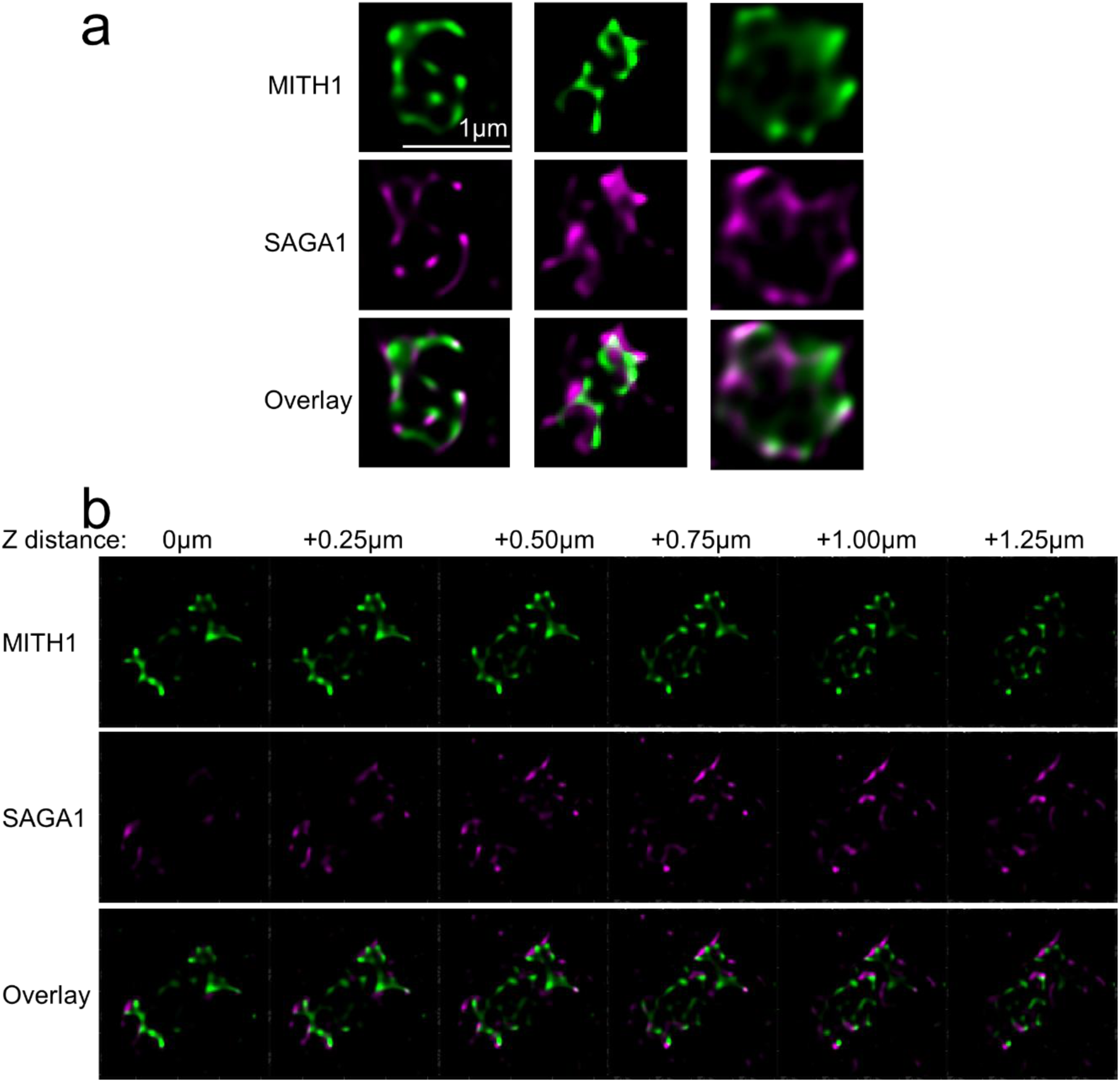
SAGA1 localizes to the exterior of MITH1, as seen by STimulated Emission Depletion (STED)-super-resolution microscopy. Each image is zoomed in on a single pyrenoid, with MITH1 fluorescence shown in green and SAGA1 fluorescence shown in magenta. **a**, Pyrenoids from three different cells are shown. **b,** Six z-planes are shown, each 0.25 µm apart from each other.

**Extended Data Figure 6.**
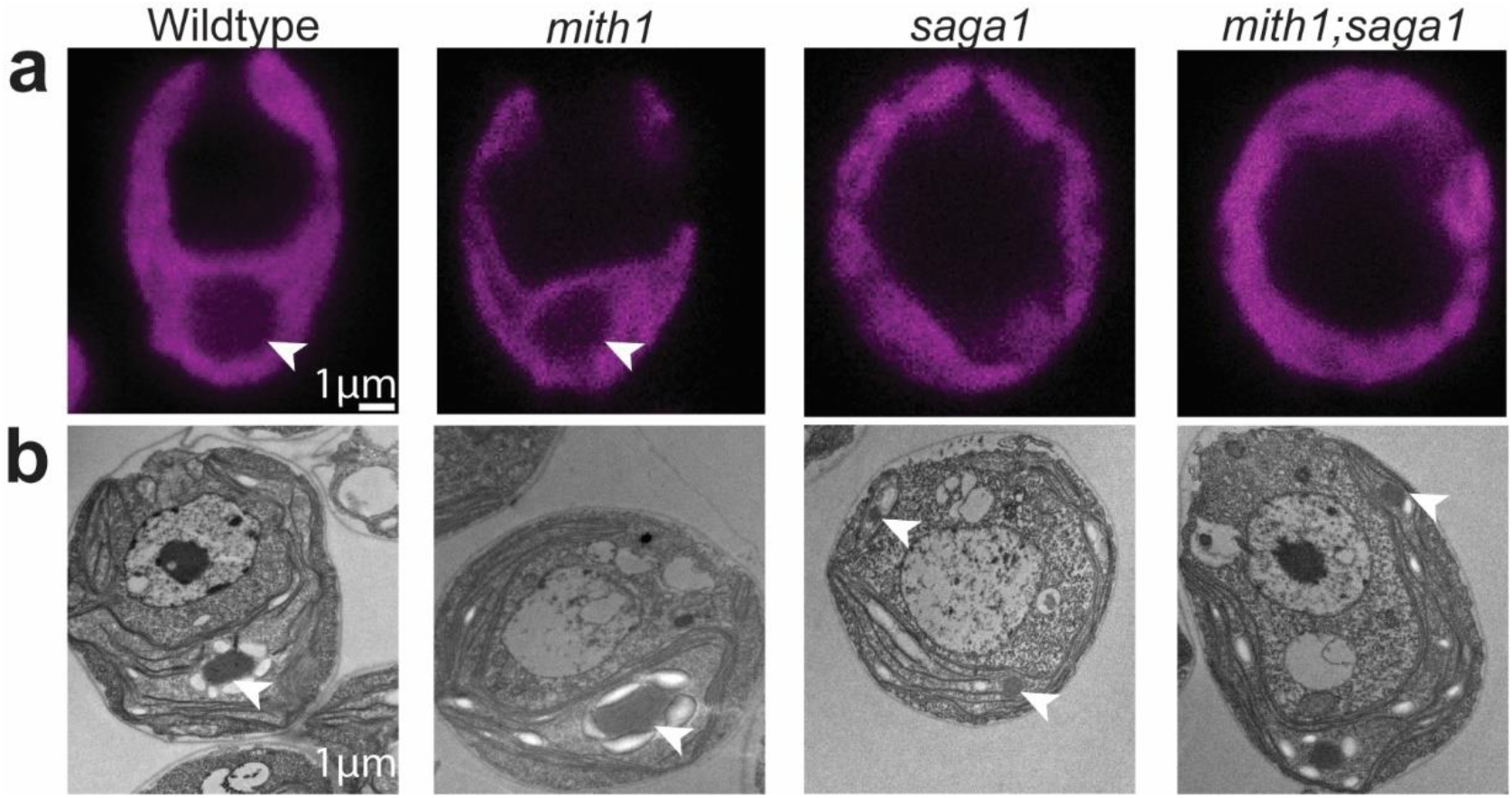
Comparisons of cell morphology between wildtype, single, and double mutants using **a**, chlorophyll autofluorescence or **b,** TEM. Arrowheads point to observed pyrenoids.

**Extended Data Fig. 7.**
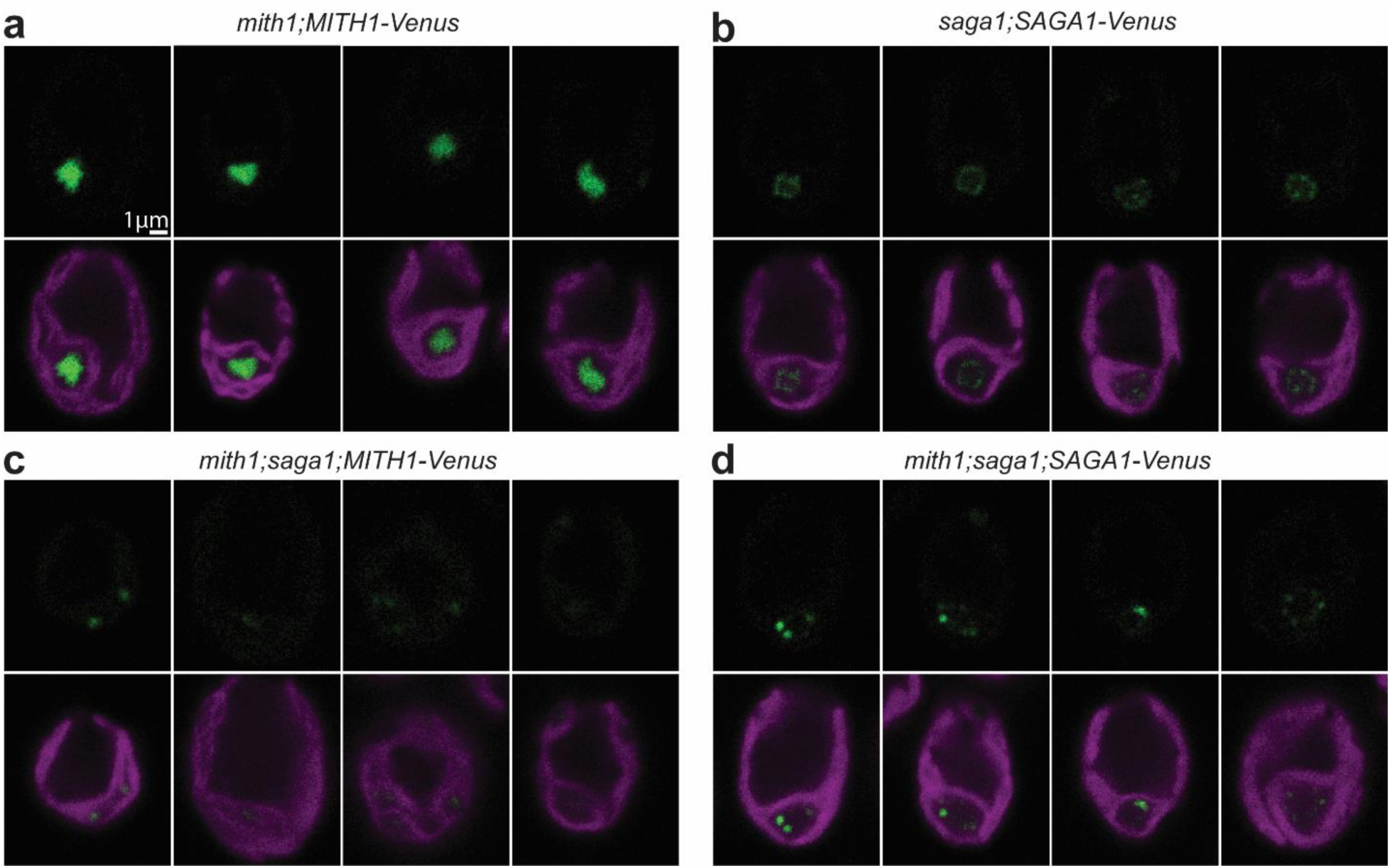
MITH1 localization is dependent on SAGA1, but SAGA1 localizes independently of MITH1. a, MITH1 in the *mith1;MITH1-Venus* complemented strain localizes to structures within the pyrenoid. **b,** SAGA1 in the *saga1;SAGA1-Venus* complemented strain localizes to puncta at the pyrenoid periphery^13^. **c**, MITH1 mis-localizes throughout the chloroplast when expressed in the *mith1;saga1* double mutant that is missing SAGA1. **d**, SAGA1 localizes to pyrenoid peripheral puncta in the absence of MITH1.

**Extended Data Fig. 8.**
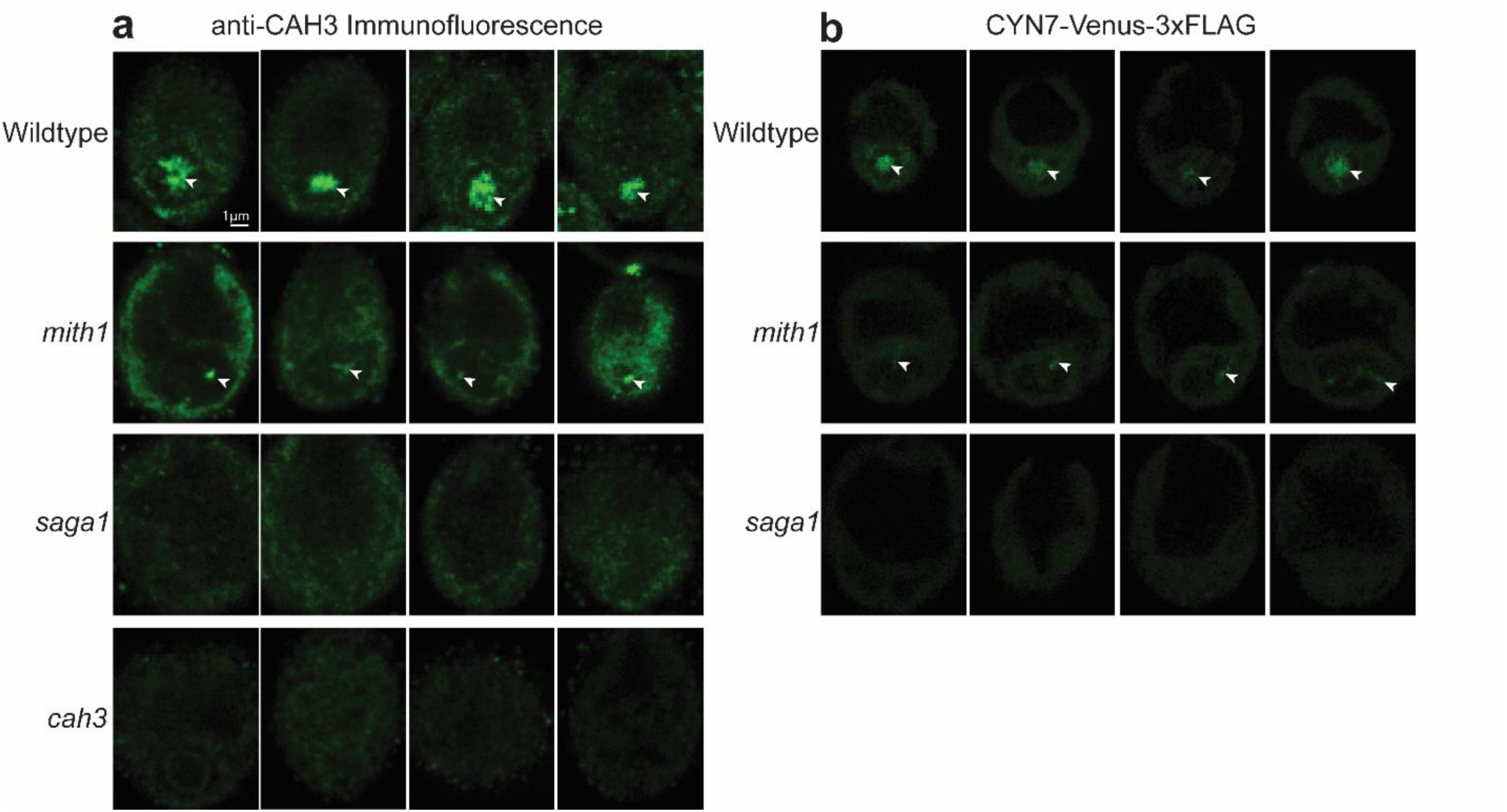
Tubule proteins CAH3 and CYN7 are mis-localized in *mith1* and *saga1* mutants. **a**, Immunofluorescence with an anti-CAH3 antibody was used to visualize localization of endogenous CAH3. Arrowheads point to bright signal corresponding to CAH3. In *mith1* mutants, CAH3 localized to small puncta and streaks at the periphery of the pyrenoid rather than to the star-like traversing network seen in wildtype cells. The *cah3* mutant was used as a negative control. **b**, Additional images of CYN7-Venus expression in live cells of the wildtype, *mith1*, and *saga1* strains. Arrows point to the wildtype tubules and *mith1* thylakoid protrusions labeled by CYN7-Venus.

**Extended Data Fig. 9.**
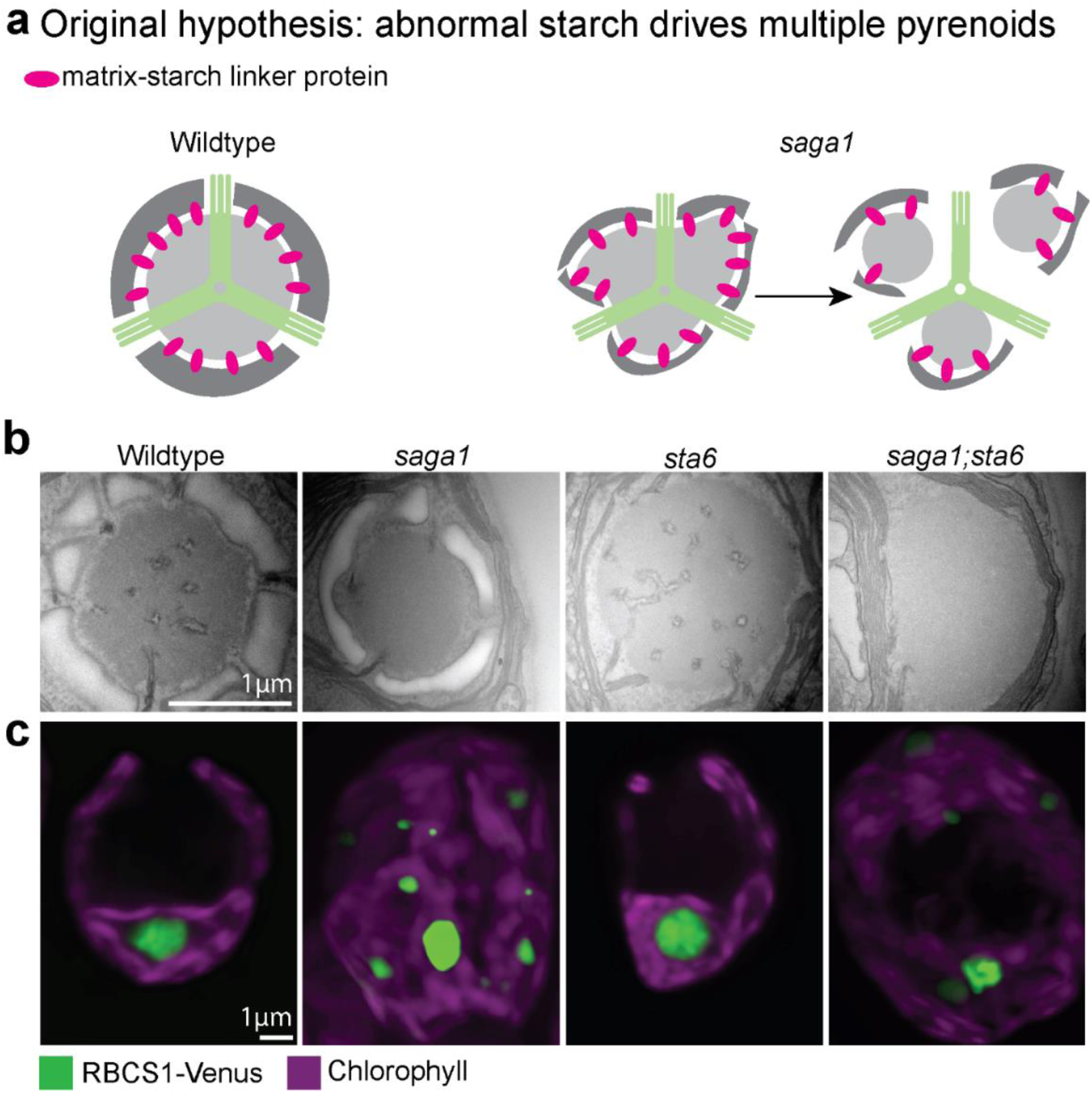
Starch is not the driver of multiple pyrenoids and missing tubules in the *saga1* mutant. **a**, In an earlier model^13^, we hypothesized that an increased surface area of misshapen starch leads to multiple pyrenoids which are missing tubules in the *saga1* mutant through interactions between a starch-matrix linker protein. **b**, TEM images of wildtype and mutant pyrenoids, including the starchless *sta6* mutant and the *saga1;sta6* double mutant. **c**, Max-z projections of RBCS1-Venus expressed in the wildtype compared to mutant backgrounds.

## Methods

### Chlamydomonas strains and culture conditions

Strains used in this study are listed in Supplementary Table 1. Cells were maintained and cultured for experiments as described previously^18^. Briefly, unless otherwise noted, cells used in imaging and immunoblot experiments were grown to mid-log phase in Tris-Phosphate (TP) liquid media in an orbital shaking incubator at room temperature (22 °C), 130 rpm, continuous cool white LED light at ∼175 µmol photons m^−2^⋅s^−1^, and in air enriched to 3 % CO_2_ to enable growth of mutants with defective CO_2_ concentrating mechanisms. 16-24 hours prior to experiments, cultures were moved to a different incubator with the same parameters except at air levels of CO_2_ to induce the CO_2_ concentrating mechanism. Cultures were harvested for experiments at a cell density of approximately 2 × 10^6^ cells/mL, as measured by a Countess II Automated Cell Counter.

**Supplementary Table 1.** List of Chlamydomonas strains used in this study and their sources.

| Chlamydomonas Resource Center ID | Strain Description | Mating type (mt) | Source | Antibiotic resistance |
| --- | --- | --- | --- | --- |
| CC-4533 | CMJ030 (CC-4533) | Minus (mt-) | Wildtype and parent strain to CLiP library<sup>27</sup> | none |
| LMJ.RY0402.133670 | *mith1* | mt- | CLiP library LMJ.RY0402.133670<sup>28</sup> | Paromomycin |
| CC-5938 | *mith1*;MITH1-Venus-3xFLAG | mt- | Transformation of *mith1* mutant with linearized MITH1-Venus-3xFLAG pRAM118 plasmid | Paromomycin, hygromycin |
| CC-5420 | *saga1* | mt- | Previously published<sup>13</sup> | Paromomycin |
| CC-5939 | CMJ030; CYN7-Venus-3xFLAG | mt- | Transformation of CMJ030 with linearized CYN7-Venus-3xFLAG pLM005 plasmid | Paromomycin |
| CC-5940 | *mith1*;CYN7-Venus-3xFLAG | mt- | Transformation of *mith1* with linearized CYN7-Venus-3xFLAG pLM005 plasmid | Paromomycin, Hygromycin |
| CC-5941 | *saga1*;CYN7-Venus-3xFLAG | mt- | Transformation of *saga1* with with linearized CYN7-Venus-3xFLAG pLM005 plasmid | Paromomycin, Hygromycin |
| CC-5617 | CMJ030;RBMP1-Venus-3xFLAG | mt- | Previously published<sup>18</sup> | Paromomycin |
| CC-5155 | CC-5155 | mt+ | Wildtype strain made through cross between CMJ030 mt-and CC-125 mt+ followed by 5 times backcrossing to CMJ030; Chlamydomonas Resource Center CC-5155 | none |
| CC-5942 | CMJ030; RBCS1-Venus-3xFLAG | Plus (mt+) | Crossing of the previously published<sup>1</sup> CMJ030;RBCS1-Venus-3xFLAG mt- strain to the wildtype CC1690 mt+ strain | Paromomycin |
|  | *Rubisco mutant D23A/E24A11* | mt- | Previously published<sup>10</sup> | none |
| CC-5943 | *saga1*;sta6 | Unknown | Crossing of *saga1* mt- with *sta6* mt+ | Paromomycin |
| CC-5944 | *saga1*;sta6;SAGA1-Venus-3xFLAG | Unknown | Transformation of *saga1*;sta6 double mutant with linearized SAGA1-Venus-3xFLAG pRAM118 plasmid | Paromomycin, Hygromycin |
| CC-5374 | *sta6* | mt+ | Chlamydomonas Resource Center C5374 | none |
| CC-5422 | *saga1*;SAGA1-Venus-3xFLAG | mt- | Previously published<sup>13</sup> | Paromomycin, Hygromycin |
| CC-5945 | *saga1*; RBCS1-Venus-3xFLAG | Unknown | Crossing of *saga1* mt- with CMJ030; RBCS1-Venus-3xFLAG mt+ | Paromomycin |
| CC-5946 | *mith1*; RBCS1-Venus-3xFLAG | Unknown | Crossing of *mith1* mt- with CMJ030; RBCS1-Venus-3xFLAG mt+ | Paromomycin |
| CC-5947 | *saga1* mt+ | mt+ | Crossing of *saga1* mt- with CC-5155 mt+ | Paromomycin |
| CC-5948 | *mith1*;saga1 | Unknown | Crossing of *mith1* mt- with *saga1* mt+ | Paromomycin |
| CC-5949 | *mith1*;saga1;MITH1-Venus-3xFLAG | Unknown | Transformation of *mith1*;saga1 with linearized MITH1-Venus-3xFLAG pRAM118 plasmid | Paromomycin, Hygromycin |
| CC-5950 | *mith1*;saga1;SAGA1-Venus-3xFLAG | Unknown | Transformation of *mith1*;saga1 with linearized SAGA1-Venus-3xFLAG pRAM118 plasmid | Paromomycin, Hygromycin |
| CC-5951 | *saga1*;RBMP1-Venus-3xFLAG | mt- | Co-transformation of *saga1* with linearized RBMP1-Venus-3xFLAG pLM164 plasmid and Hygromycin resistance gene from pRAM118 | Paromomycin, Hygromycin |
|  | *sta6*;RBCS1-Venus-3xFLAG | mt+ | Transformation of *sta6* with RBCS1-Venus-3xFLAG pRAM118 plasmid | Hygromycin |
|  | *saga1*;sta6;RBCS1-Venus-3xFLAG | unknown | Transformation of *saga1*;sta6 with RBCS1-Venus-3xFLAG pRAM118 plasmid | Paromomycin, Hygromycin |

Unless otherwise noted, the wildtype CMJ030 (CC-4533)^27^ was the background for all strains. New strains produced for this study were made through crossing and transformation, as described below. Mutants from the CLiP library were produced using insertional mutagenesis with a paromomycin resistance gene^28^.

### Arabidopsis plant material and growth conditions

Arabidopsis (*Arabidopsis thaliana*, Col-0 background) seeds were sown on compost or on half-strength Murashige and Skoog basal salt medium (Sigma) with 1 % sucrose, stratified for 3 days at 4 °C and grown at 20 °C, ambient CO_2_ and 70 % relative humidity, under 200 µmol photons m^−2^⋅s^−1^ supplied by cool white LED lights (Percival SE-41AR3cLED, CLF PlantClimatics GmbH, Wertingen, Germany) in 12 h light, 12 h dark.

### Construct design and transformation

Plasmids used for Chlamydomonas are listed in Supplementary Table 2. Open reading frames were cloned into either the pLM005^1^ (Genbank: KX077945.1) or pRAM118^28^ (Genbank: MK357711) plasmid backbones. The two plasmid backbones are identical except that pLM005 contains an AphIII cassette encoding paromomycin resistance which is replaced by the AphII cassette encoding hygromycin resistance in pRAM118. Both plasmids encode the fluorescent protein Venus followed by three copies of the FLAG epitope.

**Supplementary Table 2.** Plasmids used for transformation into Chlamydomonas.

| Plasmid | Source | *Chlamydomonas* antibiotic resistance | *E. Coli* antibiotic resistance | Restriction enzymes |
| --- | --- | --- | --- | --- |
| *MITH1-Venus-3xFLAG pRAM118* | InFusion (Takara Bio) cloning of PCR-amplified *MITH1* into HpaI-linearized PRAM118 vector in-frame with *Venus-3xFLAG* | Hygromycin | Ampicillin | EcoRV |
| *CYN7-Venus-3xFLAG pLM005* | Previously published<sup>17</sup> | Paromomycin | Ampicillin | BsaI |
| *SAGA1-Venus-3xFLAG pRAM118* | Previously published<sup>13</sup> | Hygromycin | Ampicillin | NdeI |
| *RBMP1-Venus-3xFLAG pLM164* | Previously published | Paromomycin | Ampicillin | SpeI |
| *RBCS1-Venus-3xFLAG pRAM118* | Gibson assembly (NEB) cloning of PCR-amplified *RBCS1* into HpaI-linearized pRAM118 vector in-frame with *Venus-3xFLAG* | Hygromycin | Ampicillin | NotI |

The MITH1-Venus-3xFLAG construct was made following a previously-published PCR amplification protocol^16^. A KOD-Xtreme polymerase (Takara Bio Inc, Shiga, Japan) and manufacturer’s instructions were used to amplify *MITH1* from Chlamydomonas genomic DNA. MITH1 was annealed into an HpaI-digested pRAM118 vector using the InFusion kit (Takara Bio Inc). PCR primers (ACAACAAGCCCAGTTATGTTGCGCTCTGCAAAGCC; ACCCAGATCTCCGTTGGGCGTCTTGAGAACGCTGTTGC) containing homology tails to pRAM118 were designed according to the previous protocol^16^. The plasmid’s sequence was verified by nanopore sequencing (CD Genomics) and was found to contain 6 point mutations in intron sequences; this was deemed acceptable for the purposes of mutant gene rescue experiments.

For transformation, Chlamydomonas cells were grown to mid-log phase in liquid Tris-Acetate-Phosphate (TAP) media in air with the standard settings described above. Cells were then centrifuged at 1000 g for 5 minutes and washed twice with 10 mL of MAX reagent (GeneArt MAX Efficiency transformation Reagent for Algae from Invitrogen). Cells were resuspended to a concentration of 2 x 10^8^ cells/mL in MAX reagent. For each transformation, 60 µL of cells were combined with 5 µL of DNA that had been linearized in a standard restriction digest using the enzyme indicated in Supplementary Table 2 and their manufacturer’s protocol from New England Biolabs. This mixture was incubated at 4 °C for 5 minutes and then transferred to an electrocuvette (2-mm gap, from Bulldog bio) which had been pre-cooled at 4°C. Electroporation was done using a NEPAGENE NEPA21 electroporator (parameters: Poring Pulse 250, 8, 50, 2, 40, +; Transfer Pulse: 20, 50, 50, 5, 40 +-)^27^. Immediately following electroporation, the mixture was added to 8mL TAP + 40 mM sucrose and incubated with agitation in the dark overnight. Cells were then pelleted at 600 g for 5 minutes and plated on TAP-agar solid plates with the appropriate antibiotics, either hygromycin or paromomycin. After 5 days of growth in dim light, plates were transferred to 100 µE light for 7-14 days until colonies appeared and were of a sufficient size for picking.

For constructs used to transform *Arabidopsis*, the coding sequences of SAGA1, MITH1, and EPYC1 were codon optimized for expression in land plants and cloned into binary level 2 acceptor vectors using the Plant MoClo system^29^. The 35S cauliflower mosaic virus (CaMV) promoter and cassava vein mosaic virus (CsVMV) promoter were used to drive expression.

Preliminary experiments indicated that the Chlamydomonas MITH1 protein localizes to the cytosol of *N. benthamiana*; therefore, a chloroplast transit peptide was added to MITH1 for expression in Arabidopsis. Level 2 vectors were transformed into Agrobacterium tumefaciens (AGL1) for stable insertion in Arabidopsis plants by floral dipping^30^. First, SAGA1-mCherry and EPYC1-GFP were expressed in tandem in the Arabidopsis S2_Cr_ line^26^, consisting of the *1a3b* Rubisco small subunit mutant complemented with Chlamydomonas Rubisco small subunit CrRBCS2; homozygous lines were identified in the T2 generation using the pFAST-R selection cassette^31^. Then, MITH1-mCerulean was transformed into the resulting SAGA1;EPYC1;CrRBCS2 line and transformants were selected based on their kanamycin resistance.

The indicated restriction enzyme was used to linearize the plasmid before transformation.

### Crossing of Chlamydomonas strains

Several strains used in this study were produced through crossing opposite mating-type strains following a previously published protocol^32^. Briefly, 5-6-day old cultures maintained on TAP-agar plates were transferred to N10 TAP plates, which are nitrogen-depleted, containing 10% of the nitrogen normally supplied on TAP plates. After 3 days in this nitrogen-deficient condition, cells were transferred to a 50 mL Erlenmeyer flask containing 2.5 mL sterile distilled water using a sterile loop and agitated for 2 hours on an orbital shaker in air at ∼175 µmol photons m^−2^⋅s^−1^ light. Equal volumes of the mating type + and mating type – strains were mixed and left at ∼175 µmol photons m^−2^⋅s^−1^ light without agitation for 30 minutes to 1 hour. 300 µL of the mixed cells were then plated on TAP-3 % agar plates and allowed to dry in a sterile hood. The plates were then wrapped in foil and left in the dark for 5-7 days. After this time, vegetative cells were scraped from the plate with a flame-sterilized dull scalpel or razor blade, leaving zygospores behind. Instead of dissecting tetrads, zygospores were allowed to mature into colonies and then transferred and streaked to single colonies on a new plate containing antibiotics to select for the correct genotype. For crosses to RBCS1-Venus or the starchless *sta6* mutant, further screening was done to search for fluorescence (using Typhoon scanner) or the absence of starch (tested with Lugol’s iodine treatment). The genotype was also verified by PCR.

### Transmission electron microscopy

The Chlamydomonas samples for electron microscopy were prepared at room temperature and nutated in 1 mL volumes during chemical treatments and washes unless otherwise noted. After initial centrifugation for harvesting, all pelleting was done at 3000xg for 1 minute. Approximately 50 × 10^6^ cells were harvested at 1000 g for 5 minutes and fixed in 2.5 % glutaraldehyde in TP medium for one hour. After three 5-minute washes in MilliQ water, samples were treated with a freshly prepared solution of 1% OsO_4_, 1.5% K3Fe(CN)_3_ and 2 mM CaCl_2_. After four 5-minute washes, samples were then serially dehydrated (5-minute incubations in 50%, 75%, 95% and 100% ethanol, followed by two 10-minute incubations in 100% acetonitrile). The samples were then suspended in 50% acetonitrile, 17.5% Quetol 651, 22.5% nonenyl succinic anhydride, and 10% methyl-5-norbornene-2,3-dicarboxylic anhydride and left uncapped and stationary overnight in a fume hood to allow for evaporation of the acetonitrile. The samples were then embedded in epoxy resin containing 34% Quetol 651, 44% nonenyl succinic anhydride, 20% methyl-5-norbornene-2,3-dicarboxylic anhydride and 2% catalyst dimethylbenzylamine. The resin mixture was refreshed daily for 4 subsequent days. After the last resin refresh, pellets were resuspended in 300 µL of the resin mixture and centrifuged at 30 degrees for 20 minutes at 10500 rpm in a swinging bucket rotor for microfuge tubes. They were then cured at 65 °C for 48 hours. Afterwards, ultramicrotomy was done using a DiaTome diamond knife on a Leica UCT Ultramicrotome at the Imaging and Analysis Center, Princeton University, and imaging was done on a CM100 transmission electron microscope (Philips) at 80 kV or CM200 at 200 kV.

For *Arabidopsis* TEM, leaf samples were taken from 21-d-old plants and fixed with with 4 % (v/v) paraformaldehyde, 2 % (v/v) glutaraldehyde and 0.1 M sodium cacodylate (pH 7.2). Leaf strips (1 mm wide) were vacuum infiltrated with fixative three times for 15 min, then rotated overnight at 4°C. Samples were rinsed three times with PBS (pH 7.4) then dehydrated sequentially by vacuum infiltrating with 50 %, 70 %, 80 % and 90 % ethanol (v/v) for 1 hr each, then three times with 100%. Samples were infiltrated with increasing concentrations of LR White Resin (30 %, 50 %, 70 % [w/v]) in ethanol overnight for each, then 100 % resin three times for 1 hr. The resin was polymerized in capsules at 50 °C overnight. Sections (1 μm thick) were cut on a Leica Ultracut ultramicrotome, stained with Toluidine Blue, and viewed in a light microscope to select suitable areas for investigation. Ultrathin sections (60 nm thick) were cut from selected areas and mounted onto plastic-coated copper grids. Grids were stained in 2 % (w/v) uranyl acetate then viewed in a JEOL JEM-1400 Plus TEM (JEOL, Peabody, Massachusetts, USA). Images were collected on a GATAN OneView camera (GATAN, Pleasanton, California, USA).

### Immunofluorescence

Samples for immunofluorescence were prepared according to a published protocol^33^. Briefly, cells were fixed through 20-minute treatments with 4% (w/v) formaldehyde in PBS followed by 100 % methanol at −20 °C. Cells were then washed in PBS and blocked for 1 hour in PBS containing 5 % (w/v) Bovine Serum Albumin (BSA), treated with primary antibody for 1 hour, and secondary antibody for another hour. Antibody dilutions were 1:500 in PBS for all primary and secondary antibodies except anti-CAH3 (Agrisera), which was diluted 1:200 per supplier’s recommendation.

A Rabbit polyclonal anti-CAH3 primary antibody (Agrisera) and Alexa Fluor 488-conjufated goat-anti-Rabbit secondary antibody (ThermoFisher) were used to visualize CAH3 localization. For super-resolution co-localization of MITH1 and SAGA1, the *saga1;SAGA1-Venus-3xFLAG* strain was prepared for immunofluorescence as described above; MITH1 was labeled with a Rabbit anti-MITH1 primary antibody (Yenzyme) and a STAR RED-conjugated goat anti-Rabbit secondary antibody (Abberior), and SAGA1 was labeled using a mouse monoclonal anti-FLAG primary antibody (Sigma Aldrich) and STAR ORANGE-conjugated goat anti-mouse secondary antibody (Abberior).

### Confocal microscopy

For most Chlamydomonas experiments, live cells were imaged with a Leica TCS SP5 laser scanning confocal microscope (Leica Microsystems) using a 100 x magnification oil objective, and images were processed with FIJI software (Schindelin 2012)^34^; RBCS1-Venus expressing strains were imaged using a Nikon A1R confocal laser scanning microscope, and image analysis was done on Nikon NIS Elements Software. Excitation/emission wavelengths were 514 nm/521-540 nm for Venus and 514 nm/645-725 nm for chlorophyll autofluorescence.

For the anti-CAH3 immunofluorescence experiment, slides were imaged on a Leica TCS SP5 laser scanning confocal microscope (Leica Microsystems). Excitation/emission wavelengths were 488 nm/515-535 nm. For the super-resolution co-localization of MITH1 and SAGA1, imaging of MITH1 and SAGA1-Venus-3XFLAG was performed using a Nikon A1R confocal system with an attached Abberior STEDYCON STimulated Emission Depletion (STED) unit and a 100x magnification/1.45 NA oil immersion objective. Super-resolution was achieved with a 775 nm pulsed depletion laser. The STAR RED fluor (corresponding to MITH1) was excited with a 640 nm laser and its emission was captured with a 660 nm detector. The STAR ORANGE fluor (corresponding to SAGA1-Venus-3xFLAG) was excited with a 561 nm and its emission was captured with a 616 nm detector. Image acquisition was performed using the Abberior STEDYCON browser-based application. After acquisition, Nikon NIS Elements software was used to optimize image quality using the Denoise.ai and 3D Deconvolution tools.

*Arabidopsis* leaves were imaged with a Leica TCS SP8 laser scanning confocal microscope (Leica Microsystems). Excitation/emission wavelengths were 488 nm/502–532 nm for GFP, 405nm/462-487 for mCerulean, 552nm/599-619 for mCherry and 488 nm/680–750 nm for chlorophyll autofluorescence. Processing of these images was done with Leica LAS X software.

### Analysis of RBCS1-Venus puncta

To determine the number of condensates in each cell, Nikon NIS Elements software was used to threshold images based on Venus signal intensity, disregarding parts of the image with no or very low intensity while represent continuous clusters of high-intensity pixels as binary objects each representing a distinct RBCS1 condensate. Object identity was conserved over multiple z-slices, meaning that each was correctly identified as a single condensate even if it appeared across numerous slices.

Nikon NIS Elements software was used to measure the size of each condensate. For each condensate, which was detected and turned into a binary object as described above, the cross-sectional area was measured at each z-slice it appears in, and the largest one was reported as the representative area of the condensate, as condensates are approximately spherical. This provided a reliable way of comparing condensate size in lieu of measuring volume, which is complicated by limits to z-resolution. To calculate the proportion of RBCS1 signal contained in the largest condensate of a given cell, a ratio was taken between the largest condensate area in the cell and the sum of all condensate areas in the cell.

Data tabulation, statistical analysis, and data visualization were each performed using the GraphPad Prism 9 software. A two-tailed Mann-Whitney U test was used to test for significance between strains when comparing the proportion of rubisco fluorescence in the largest condensate of each cell.

### Western blot

For Chlamdyomonas experiments, the PVDF membrane (Millipore) was blocked for 1 hour in 5 % milk in TBST (Tris buffered Saline with 0.1% tween-20), then incubated with primary antibody for 1 hour. After 3 × 10 minute washes in TBST, the blot was treated with secondary antibody for 1 hour and washed 3 × 10 minutes again. The secondary antibody was diluted 1:10,000. Anti-SAGA1 (Yenzyme) and anti-MITH1 (Yenzyme) were diluted 1:1000. Anti-RBMP1 (gift from L. Mackinder) and anti-CAH3 (Rabbit polyclonal, Agrisera) were diluted 1:2000. Anti-tubulin (mouse monoclonal, Sigma-Aldrich) was diluted 1:4000. Blots were labeled using an enhanced chemiluminescence system (Advansta WesternBright ECL kit) and imaged on an iBright Imaging System (ThermoFisher).

Protein analyses for *Arabidopsis* were done as follows. Soluble protein was extracted from frozen leaf material of 21-d-old plants in protein extraction buffer (20 mM Tris-HCl pH 7.5 with 5 mM MgCl2, 300 mM NaCl, 5 mM DTT, 1 % Triton X-100 and cOmplete Mini EDTA-free Protease Inhibitor Cocktail (Roche)). Samples were heated at 80 °C for 15 min with 1 x Bolt LDS sample buffer (ThermoFisher Scientific) and 200 mM DTT. Extracts were centrifuged and the supernatants subjected to SDS-PAGE on a 12 % (w/v) polyacrylamide gel and transferred to a nitrocellulose membrane. Membranes were probed with rabbit serum raised against SAGA1 (1:1,000) or EPYC1 (1:2,000), or with mouse anti-Actin (Agrisera, AS21 4615) or mouse anti-GFP (Santa Cruz, sc-9996), followed by HRP-linked goat anti-rabbit IgG (Abcam) at 1:10,000 dilution, or rabbit anti-mouse IgG (Agrisera) at 1:10,000 dilution. Membranes were visualized using Pierce ECL Western Blotting Substrate (Life Technologies).

### Spot tests

Cells were grown to mid-log phase in liquid TAP medium. They were then pelleted at 1,000 g for 5 minutes and resuspended to a concentration of 6 × 10^5^ cells/mL in Tris-phosphate (TP) medium. 10µL of each sample were spotted onto TP-agar plates. Once the spots were dry, plates were then placed under 100 µmol photons m^−2^⋅s^−1^ of light in variable CO_2_ levels which included 40 parts per million (ppm) CO2, ambient CO2 (approx. 415 ppm), or 3% CO_2_ for 7 days before imaging on a Phenobooth imager. For the 40 ppm condition, plates were first kept in ambient air for the first 24 hours before being placed in a 40 ppm chamber for the remaining 6 days.

### Cell fractionation protocol

Strains were grown in TAP medium at air CO_2_ until they reached 2×10^6^ cells/mL density. The D23A/E24A11 mutant is sensitive to light and was therefore grown at low light (20 µmol photons m^−2^⋅s^−1^). Cells were centrifuged for 5 minutes at 1,500 g and the pellet was weighed and resuspended in 2x volume of lysis buffer (50mM HEPES, 10mM KOAc, 2mM Mg(OAC)2, 1mM CaCl2, pH 7.0 + protease inhibitors). Cells were sonicated in a probe tip sonicator (Qsonica) on ice for 5 minutes, 3 second pulse, 60% amplitude. The lysate was centrifuged at 2,000 g for 10 minutes to remove intact cells, and 50 μL of supernatant was collected as the whole cell lysate (WCL) sample. Unless otherwise indicated, the remaining supernatant was centrifuged at 15,000 g for 30 minutes and pellet and supernatant were collected for analysis. Laemmeli sample buffer was added to samples followed by boiling at 95°C for 10 minutes. Cell lysates were separated on an SDS-polyacrylamide gel (BioRad) and transferred to a PVDF membrane using a semi-dry system (BioRad Trans-Blot SD Cell). Antibody dilutions were as indicated above. Primary antibody was added overnight at 4°C, followed by three washes in 1x TBS-0.1% Tween-20. Secondary antibody was added for 1 hour at room temperature, followed by three additional washes in TBS-T. Blots were labeled using an enhanced chemiluminescence system (Advansta WesternBright ECL kit) and imaged on an iBright Imaging System (ThermoFisher).

## Acknowledgements

We thank members of the Jonikas laboratory as well as N. Wingreen and C. Brangwynne for discussions and feedback on the manuscript; J. Whitney for assistance in figure preparation; G. Riddihough and M. Bao for editing the manuscript; the Princeton University Confocal Microscopy core facility, a Nikon Center of Excellence, and manager G. Laevsky for fluorescence microscopy support; the Princeton Imaging and Analysis Center director N. Yao and senior research specialists J. Schreiber and P. Shao, and Stephen Mitchell of the Edinburgh School of Biological Sciences TEM facility for electron microscopy support; R. Patel at the Rutgers University Core Imaging Lab for preliminary ultramicrotomy services; M. Wühr and L. Martin for their help with acquiring mass spectrometry data; S. He for providing the *CMJ030;RBMP1-Venus mt+* strain; and L. Mackinder for discussions and for generously providing the anti-RBMP1 antibody. This project was supported by grants from the U.S. National Institutes of Health (T32GM007388), Howard Hughes Medical Institute and Simons Foundation Faculty Scholar program (55108535), U.S. National Science Foundation (MCB-1935444), U.S. National Institutes of Health (1R01GM140032-01), U.S. Department of Energy (DE-SC0020195), and Howard Hughes Medical Institute Investigator program to M.J.; and UK Research and Innovation Biotechnology and Biological Sciences Research Council (BB/S015531/1) and Leverhulme Trust (RPG-2017-402) to A.M. *Arabidopsis* TEM was carried out with the support of the Wellcome Trust Multi - User Equipment Grant (WT104915MA).

## Author contributions statement

J.H.H. performed Chlamydomonas TEM experiments and generated new Chlamydomonas strains used in this paper; N.A. performed all *Arabidopsis* experiments; J.H.H and A.K.B. prepared samples for immunofluorescence and performed confocal microscopy; A.K.B. performed statistical analysis; J.H.H. and S.E. performed cell fractionation and Western blot; J.H.H. and E.F. validated Chlamydomonas TEM experiments; L.W. provided early access to CYN7 protein localization information; M.K. provided early access to mutant proteomics data; F.F., J.V.B., and R.E.J. provided early access to mutant pooled screen data; J.H.H., N.A., A.J.M., and M.C.J. designed experiments and analyzed and interpreted the data; and J.H.H. and M.C.J. wrote the manuscript, with input from others.

